# Biobank-Scale Polygenic Prediction in Admixed Populations Using Local Ancestry via the Group Lasso

**DOI:** 10.64898/2026.03.14.711808

**Authors:** David Bonet, James Yang, Trevor Hastie, Alexander G. Ioannidis

## Abstract

Polygenic risk models trained in one ancestry often fail to perform well in others, in part due to linkage disequilibrium and allele frequency differences across ancestries. In response, separate models trained for specific ancestries have been introduced. However, this single ancestry approach is untenable in admixed groups, which have ancestry from multiple sources varying across the genome and between individuals. Here we present Combine, a biobank-scale sparse regression framework that augments per-variant genotype features with local ancestry dosages and fits all effects jointly using a variant-level group lasso penalty. In 99,298 admixed participants from the All of Us Research Program, Combine substantially outperforms state-of-the-art multi-ancestry summary-statistic approaches (e.g., 144% relative improvement over PRS-CSx for white blood cell count). Furthermore, it matches or improves upon the predictive performance of highly optimized individual-level models (iPGS/snpnet) across seven of nine evaluated phenotypes, while uniquely providing locus-level interpretability to disentangle shared allelic effects from ancestry-linked tagging. An ancestry-specific extension, Combine-S, estimates haplotypic ancestry-associated SNP effects together with local-ancestry terms, enabling systematic identification of ancestry-dependent effect magnitudes and sign differences at established and plausible loci. Finally, we show that incorporating external GWAS evidence through group-specific penalty weights improves LDL cholesterol prediction without pre-filtering variants. Together, Combine provides a scalable framework for polygenic modeling that prioritizes efficient local ancestry-aware modeling and interpretation in admixed biobanks.

## Introduction

Biobanks linking genomic variation to complex traits and disease risk now enable genetic analyses of unprecedented scale, particularly genome-wide association studies (GWAS) and polygenic risk score (PRS) construction [1, 2]. However, as most cohorts remain predominantly of European-descent [3, 4], while heterogeneity across genetic ancestries often limits the portability of a PRS model outside the ancestry from which it was constructed, a gap is developing in model accuracy for non-European populations [5–10]. Moreover, performance of existing PRS models in admixed individuals is particularly poor [11, 12]. Addressing this gap requires models that move beyond a single effect per variant framework or adjustments based only on aggregate ancestry proportions, but that can instead account for ancestry at the relevant locus at a genomic window level resolution.

Admixture is a term for understanding genomes as a mosaic of chromosomal segments (haplotypes) deriving from different genetic ancestries (Figure 1A,B). Local ancestry inference (LAI) methods can label the ancestry of each of these haplotype segments along the chromosome with high accuracy [13, 14]. These segments can have divergent allele frequencies and linkage disequilibrium (LD) patterns depending on their genetic ancestry, which will differ for a given segment even between individuals in the same family, due to stochastic recombination and independent assortment during meiosis. This makes a single population-wide model for such cohorts challenging. Because PRS often rely on tested SNPs that are correlated with a disease, but that are not necessarily the causal variants [2], and because correlations of neighboring variants (i.e., LD) differ amongst haplotypes of different ancestries, the effect size of tagging SNPs can vary widely depending on the genetic ancestry of the haplotype surrounding them [15]. As a result of this phenomenon alone, PRS accuracy can vary along the genome of an admixed individual, depending on their personal mosaic of ancestry blocks, and this can produce directional bias in standard PRS for admixed individuals [16].

**Figure 1.**
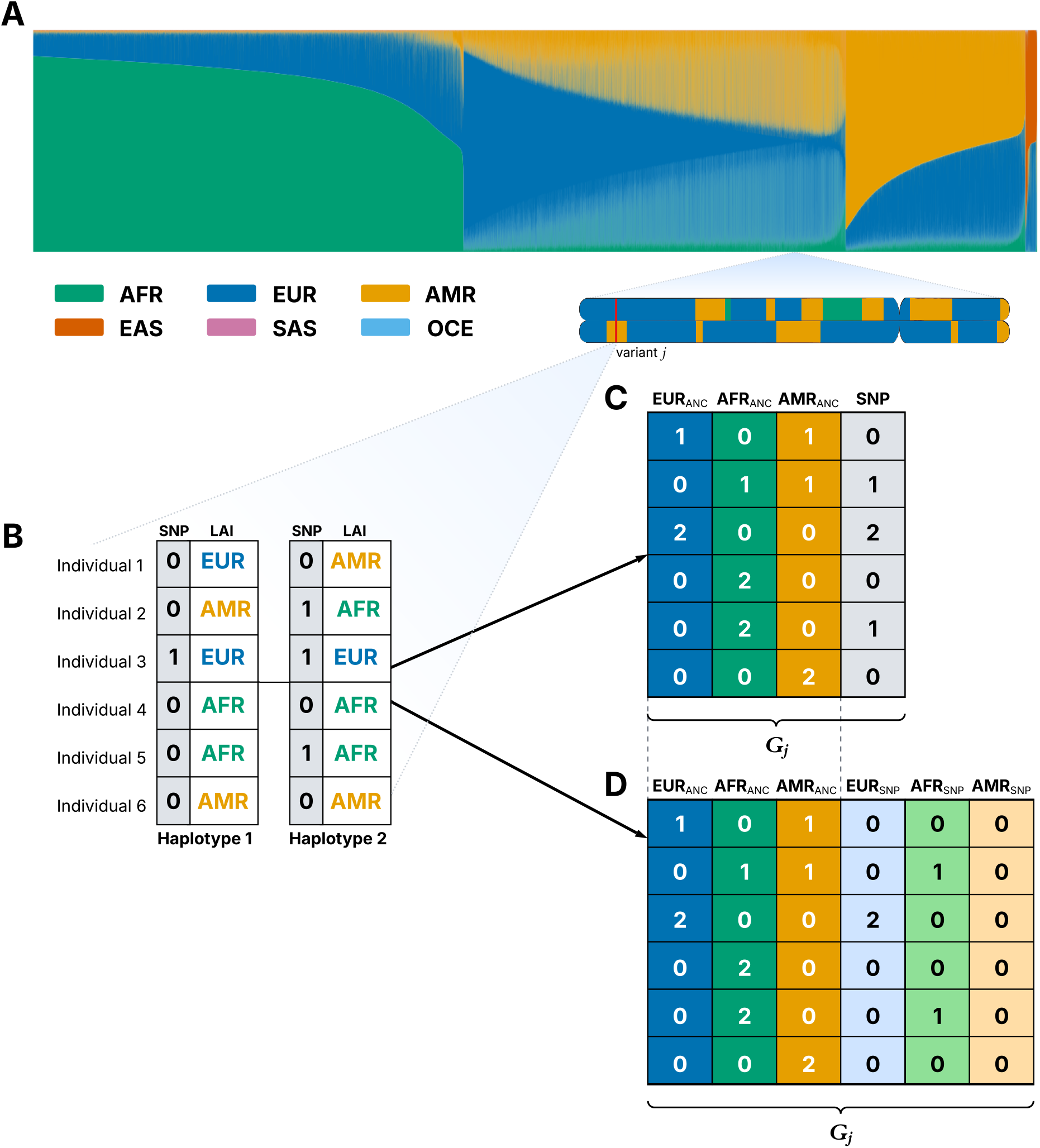
Combine overview. (**A**) Genetic ancestry proportions across 99,298 admixed participants, and chromosome painting example from local ancestry inference (LAI) in 99,298 admixed All of Us participants. (**B**) Schematic locus showing SNP genotypes (0=ref, 1=alt) and genetic ancestry (LAI) for 6 individuals. (**C**) Combine-R: unphased allele dosage plus LAI per variant. (**D**) Combine-S: ancestry-specific allele dosages plus LAI (see Methods for further model specifications).

In addition, admixture mapping has repeatedly identified genetic ancestry-enriched loci in self-identifying African American and Hispanic/Latino population cohorts, for example the Duffy antigen receptor locus for white blood cell count and the 8q24 region for prostate cancer, underscoring the value of local ancestry for trait mapping [17].

Meanwhile genetic ancestry has been shown in some cases to track disease biology, including higher rates of somatic mutation patterns in cancer [18–21]. Given that populations with substantial admixture constitute a large and growing fraction of the world population [22], there is a pressing need for prediction models that jointly leverage genotype and local ancestry along the genome, rather than averaging effects across ancestries or relying on genome-wide (global) ancestry adjustments.

Methodologically, the field has begun to move beyond univariate GWAS toward high-dimensional sparse regression that estimates many effects simultaneously. Scalable lasso regularization enables simultaneous variable selection and estimation in ultra high-dimensional settings [23]. Methods that scale to biobanks such as snpnet [24–26] and include admixed individuals [27] often outperform summary-statistics PRS, yet they still average ancestry effects into a single coefficient per variant.

Complementary to individual-level sparse regression, multi-ancestry summary-statistic methods such as PRS-CSx improve transfer across single-ancestry cohorts by using genetic ancestry-matched LD reference panels [28]. However, these panels assume a single ancestry background per LD reference, making them theoretically ill-suited in the case of an admixed individual’s genome, which carries multiple genetic ancestries along its length. To quantify this limitation, we directly benchmark against PRS-CSx to evaluate how individual-level local ancestry modeling compares to summary-statistic approaches in highly admixed cohorts.

Beyond transfer methods, local-ancestry-aware association approaches, such as Tractor [29], have demonstrated increased power and the ability to detect ancestry-enriched novel associations, but inherit univariate limitations for prediction. While recent methods like GAUDI [30], HAUDI [31], and SDPR_admix [32] have begun to integrate local ancestry into prediction, they rely on summary statistics and pre-selection steps, or are currently implemented for only two-way or three-way admixture. Crucially, these approaches cannot be feasibly applied to full biobank genotype matrices of this scale (*N* ≈ 100,000) with high-dimensional ancestry vectors using standard compute resources.

Prior All of Us PRS work grouped participants by continental genetic ancestry using an averaging approach across their entire genome, namely PCA-based projections and a supervised classifier, with additional analyses using genome-wide, averaged genetic ancestry as a continuum. Such genome-wide averaging approaches completely ignore the mosaic of haplotype variation from divergent sources along the genome of admixed individuals, which varies dramatically even between full siblings in the same family, and can be modeled by local ancestry [33]. Our study focuses on such admixed individuals and trains predictors that jointly use SNPs and per-individual, haplotype-level local ancestry, instead of genome-wide global ancestry averaging that collapses this information. To address this, we introduce Combine, a variant-level grouped sparse regression framework for admixed genomes. Combine expands each SNP into a small set of genotype and local-ancestry features and fits all effects jointly using a group lasso penalty, performing selection at the locus level while retaining interpretable within-locus decompositions. Our design captures within-genome ancestry heterogeneity of an individual’s specific haplotypes, which global (genome-wide) ancestry averaging approaches cannot. This framework makes local ancestry usable for end-to-end prediction without ancestry-binned training, ancestry-matched LD matrices, or summary-statistic pre-filtering, and produces coefficients that can be inspected to distinguish shared allelic effects from ancestry-linked tagging or regional haplotype structure. Algorithmically, Combine is engineered for fast biobank-scale training with millions of predictors and hundreds of thousands of samples, using efficient implementation strategies that support any number of ancestries. We further allow and propose incorporation of external GWAS evidence through penalty weights, guiding sparsity toward well-supported loci while keeping all variants eligible in the individual-level fit.

We evaluate Combine on 99,298 individuals with substantial admixture from the All of Us Research Program on nine binary and quantitative phenotypes. We test the hypotheses that jointly modeling genotype and local ancestry with group sparsity improves predictive accuracy for admixed individuals, enhances portability relative to ancestry-agnostic models, recovers known signals, and reveals biologically meaningful haplotypic ancestry-associated effects that are obscured by averaging. We further evaluate a prior-weighted version of Combine that incorporates external GWAS summary statistics through the penalty, without prefiltering features, and show that it improves prediction for LDL cholesterol in admixed cohorts. Together, these results point to a practical path for equitable, ancestry-aware polygenic prediction in admixed populations.

Our contributions are threefold. First, we propose two ancestry-aware grouped designs, Combine-R and Combine-S (Figure 1C,D), that integrate local ancestry inference outputs directly into sparse regression for polygenic prediction. Second, we provide an end-to-end implementation that scales these ancestry-expanded designs to biobank matrices by using matrix-free optimization and compressed genotype and ancestry encodings. Third, we empirically evaluate accuracy, sparsity, and coefficient interpretability in a large admixed cohort, including an extension that incorporates external GWAS evidence through weighted group penalties.

## Results

### Combine-R improves prediction in admixed participants

We benchmark Combine against a SNP-only individual-level model fit with `snpnet` (iPGS) and the multi-ancestry summary-statistic method PRS-CSx. We evaluate all methods across nine phenotypes in 99,298 admixed All of Us participants, using identical train, validation, and test splits and the same covariates. For quantitative traits we report *R*^2^ on the test set after covariate adjustment. For binary traits we report test AUC. When compared to the state-of-the-art multi-ancestry summary-statistic approach (PRS-CSx), Combine-R yielded substantial improvements, achieving a maximum relative improvement of 144.2% for white blood cell count (WBC), 73.1% for platelet count, and 25.0% for C-reactive protein (CRP). Furthermore, when compared to the highly optimized, individual-level SNP-only baseline (`snpnet`), Combine-R successfully matched or improved predictive performance on seven out of nine phenotypes, achieving an improvement of 4.2% for CRP. While `snpnet` yielded a slightly higher accuracy for LDL cholesterol without priors, incorporating external GWAS priors into Combine-R resulted in a 4.1% improvement over the `snpnet` baseline. Beyond prediction, Combine-R yields an interpretable per-locus decomposition: each selected variant has a SNP effect and local-ancestry effects that quantify the haplotypic ancestry background modulates the variant’s contribution. This could arise through differences in LD or in the prevalence of different causal variants across ancestry haplotypes. This supports locus-level auditing and hypothesis generation in settings where standard SNP-only models conflate allelic effects with ancestry-linked tagging. Genome-wide coefficient maps make this decomposition explicit (Figure 3A). For log(WBC), the Combine-R local-ancestry terms show a sparse set of nonzero ancestry effects, dominated by the expected signal at the Duffy locus (*ACKR1* /rs2814778) on Chromosome 1.

We report numeric comparisons in Table 1. Predictive performance across methods with 5-fold cross-validation for representative phenotypes, together with computational timing bench-marks, is shown in Figure 2.

**Table 1.** Predictive performance on the held-out admixed test set. Values are *R*^2^ for quantitative traits and AUC for binary traits. Bold indicates the best-performing method for each phenotype. Trait type is indicated as quantitative or binary. Combine-S is included primarily for interpretability (ancestry-specific SNP effects, see below). Uncertainty across folds is shown as standard errors in Figure 2 and Figure S1.

| Phenotype | Type | PRS-CSx | iPGS(snpnet) | Combine-R | Combine-S |
| --- | --- | --- | --- | --- | --- |
| Log-transformed white blood cell (WBC) count | Quantitative | 0.043 | 0.103 | **0.105** | 0.104 |
| C-reactive protein (CRP) | Quantitative | 0.020 | 0.024 | **0.025** | 0.021 |
| Platelet count | Quantitative | 0.026 | 0.044 | **0.045** | 0.038 |
| High-density lipoprotein (HDL) cholesterol | Quantitative | 0.077 | 0.136 | **0.138** | 0.129 |
| Low-density lipoprotein (LDL) cholesterol | Quantitative | 0.011 | **0.061** | 0.056 | 0.049 |
| Triglycerides | Quantitative | 0.044 | **0.070** | 0.067 | 0.062 |
| Estimated glomerular filtration rate (eGFR) | Quantitative | 0.070 | **0.084** | **0.084** | 0.082 |
| Chronic kidney disease (CKD) | Binary | 0.738 | **0.739** | **0.739** | 0.738 |
| Colorectal cancer | Binary | 0.743 | **0.753** | **0.753** | 0.752 |

**Figure 2.**
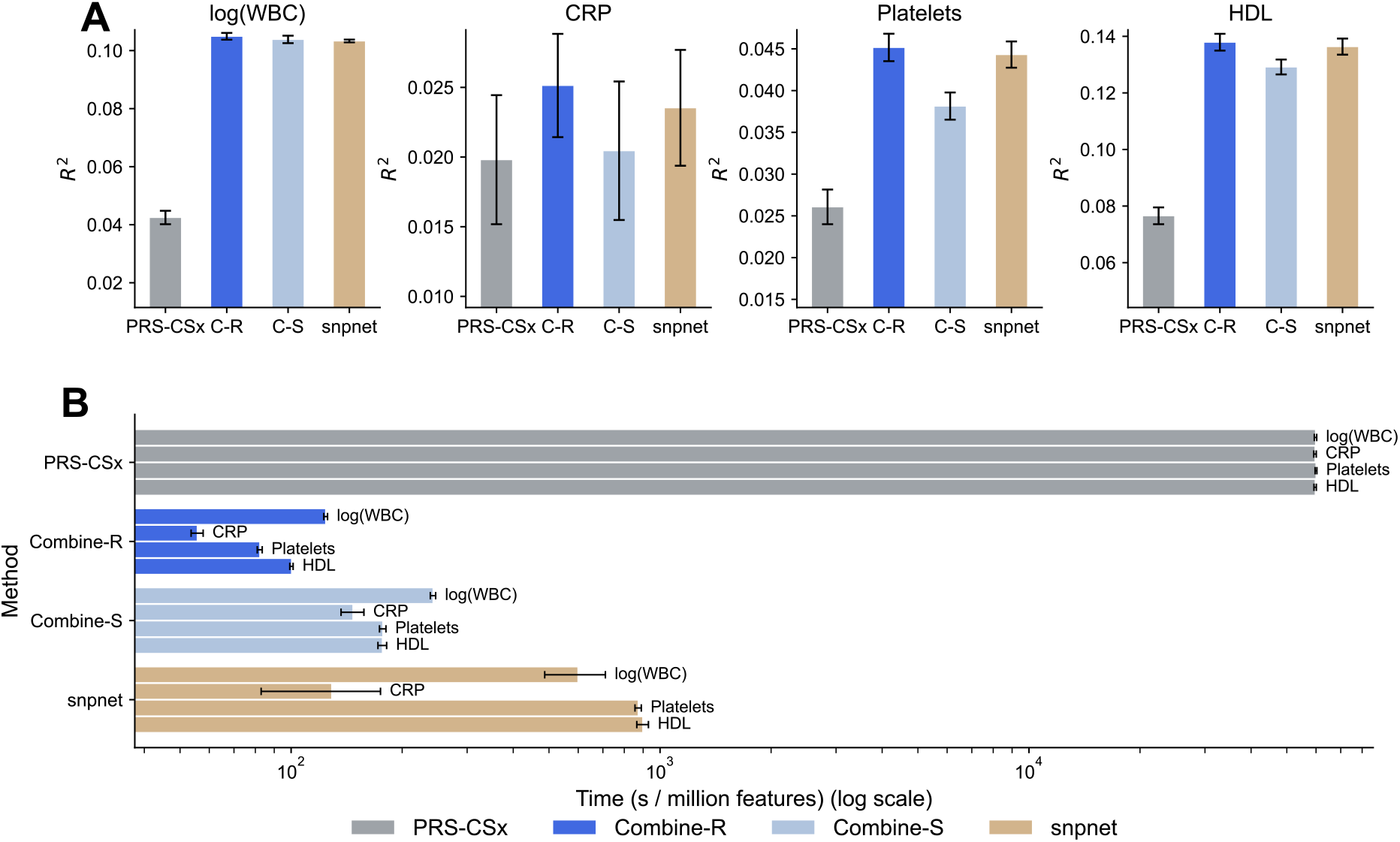
Method comparison and computational efficiency. (**A**) Mean predictive performance (*R*^2^) across 5-fold cross-validation for log(WBC), CRP, platelets, and HDL cholesterol, comparing PRS-CSx, Combine-R, Combine-S, and `snpnet` (iPGS). (**B**) Mean wall-clock training time per million features (seconds per million features). Bars show fold means and error bars indicate the standard error of the fold means (computed across the 5 cross-validation folds). Combine-R and Combine-S are significantly more efficient, leveraging compressed matrix operations to handle the expanded feature space without a proportional increase in runtime.

### Computational efficiency and scalability

We evaluated the computational cost of the different methods by comparing the mean training time per cross-validation fold normalized by the number of features processed (Figure 2B). Combine demonstrates superior efficiency, with per-feature training times 3 to 6 times faster than `snpnet`. This efficiency is driven by the solver, which utilizes compressed genotype-ancestry block operations to minimize memory traffic and accelerate gradient computations, typically completing genome-wide fits in under 20 minutes per fold. These results confirm that Combine scales to biobank-level data (*N* ≈ 100,000) with complex ancestry models without requiring specialized high-performance computing clusters beyond standard high-memory nodes.

### Ancestry-dependent effects at known and plausible loci

We examined per-ancestry SNP and local ancestry coefficients from Combine-S at selected loci to illustrate the model’s capacity to capture diverse genetic architectures. Table 2 summarizes these coefficients.

**Table 2.** Ancestry-associated SNP and local ancestry coefficients of selected variants from Combine-S. Subscripts denote whether the coefficient corresponds to the ancestry-specific allele dosage (SNP) or the local ancestry dosage (ANC). * Indicates a variant selected by the model that tags the specified neighboring gene.

| rsID | Region | CHR:BP:A1:A2 | EUR<sub>SNP</sub> | AFR<sub>SNP</sub> | AMR<sub>SNP</sub> | EUR<sub>ANC</sub> | AFR<sub>ANC</sub> | AMR<sub>ANC</sub> | Phenotype |
| --- | --- | --- | --- | --- | --- | --- | --- | --- | --- |
| rs10908722 | *ACKR1* * | 1:159439944:G:T | 0.003 | -0.008 | 0.002 | 0.003 | -0.007 | 0.002 | log(WBC) |
| rs34065661 | *CETP* | 16:56962023:C:G | 0.052 | 1.432 | -0.004 | -0.030 | 0.059 | 0.009 | HDL |
| rs4783961 | *CETP* | 16:56960982:G:A | 0.117 | 0.780 | -0.007 | -0.020 | 0.046 | -0.005 | HDL |
| rs5883 | *CETP* | 16:56973441:C:T | 0.445 | 0.906 | 0.009 | 0.048 | 0.046 | -0.033 | HDL |
| rs738409 | *PNPLA3* | 22:43928847:C:G | -1.449 | -0.767 | -0.651 | 0.096 | 0.201 | -0.376 | Platelet count |
| rs6887079 | *TENM2* | 5:167439431:C:T | -0.014 | 0.011 | -0.002 | -0.005 | -0.004 | 0.009 | Chronic kidney disease |
| rs1885722 | *AKAP6* | 14:32367316:T:G | -0.010 | 0.015 | 0.003 | -0.011 | 0.009 | 0.003 | Colorectal cancer |

### Replication of known biology and differential penetrance

We first validated our model against loci with established ancestry-associated architectures. The variant with the largest effects for log(WBC) is rs2814778 in Combine-R, and is a variant linked to it in Combine-S. This SNP has a classic effect at the *ACKR1* promoter, the Duffy-null genotype CC, which is at much higher frequency in African ancestry and results in lower neutrophil counts [34]. African haplotype associated effects are negative and large in magnitude, compared to opposite and smaller effects in the other genetic ancestry haplotypes. Notably, Combine-S selects this tagging SNP rather than the population-specific causal variant due to the mechanics of the group lasso penalty and the recessive nature of the Duffy-null phenotype. Because the Duffy-null allele is nearly absent in non-African haplotypes and its recessive effect violates strict linear additivity, its non-African columns lack variance, risking the entire variant group being shrunk to zero. By selecting a tagging SNP, the model captures the strong negative Duffy-null effect in African haplotypes via LD, while simultaneously finding other non-zero variance (and opposing effects) in haplotypes of other ancestries. All of Us phenome-wide analyses show ancestry-associated patterns at this locus [35], and Van Driest et al. [36] and Chung et al. [37] discuss clinical implications of its screening in patients with African-like ancestry. Similarly, in the *CETP* region (chr16), we correctly recovered known ancestry divergence in HDL cholesterol associations. The three variants with the largest divergence between European and African haplotypes for their SNP effect sizes (|EUR_SNP_ − AFR_SNP_|) are all located in or near *CETP*: rs34065661, rs4783961, and rs5883. This is consistent with prior work reporting strong genetic ancestry-dependent associations with HDL cholesterol in this region. For example, Pirim et al. [38] and Chang et al. [39] identified significant associations for these variants in African or African American populations that were attenuated or absent in non-Hispanic Whites, or vice versa. Combine-S correctly recovers these known ancestry-associated signals with effect directions and magnitudes consistent with the literature.

We also captured differential haplotypic ancestry-associated phenotypic penetrance at *PNPLA3* rs738409, a causal variant for nonalcoholic fatty liver disease and increased liver fat risk [40]. We observed negative SNP coefficients for platelet count across haplotypic ancestries, a finding that is biologically plausible given that thrombocytopenia is a known downstream consequence of liver pathology. Notably, the SNP effect size was nearly twice as strong in European-like haplotypes (EUR_SNP_ = −1.449) compared to African-like haplotypes (AFR_SNP_ = −0.767). This aligns with Cox et al. [41], who note that while rs738409 is a risk locus in African American cohorts, this population exhibits lower overall susceptibility to hepatic steatosis than European Americans. While this reduced susceptibility is clinically attributed to population-level differ-ences in fat partitioning and lipid metabolism [42–44], our attenuated coefficient in the African haplotype demonstrates that this lower penetrance also has an effect specific to this variant, as global ancestry covariates already adjust for baseline phenotypic differences and environment associations associated with overall ancestry proportions.

### Opposing regulatory effects in kidney and cancer

A key advantage of ancestry-specific haplotypic modeling is the ability to detect “sign flips,” extreme cases where a variant is associated with protection in one haplotypic ancestry but associated with risk in another. We observed this striking pattern across several loci related to chronic kidney disease (CKD) and cancer.

#### *TENM2* rs6887079 (chr5:167439431:C:T)

This variant is in the teneurin transmembrane protein 2 (*TENM2*) gene. *TENM2* is expressed in the kidney, specifically in tubular cells, consistent with an epithelial signaling role [45], and its expression correlates with higher estimated glomerular filtration rate (eGFR) and reduced renal fibrosis. In a large cohort of individuals of European ancestry with diabetes, Sandholm et al. [46] linked an intronic *TENM2* variant to lower risk of CKD and diabetic kidney disease. Our results recapitulate this protective effect in European-like ancestry haplotypes (EUR_SNP_ = −0.014), but identify an opposite, risk-increasing effect in African-like ancestry haplotypes (AFR_SNP_ = 0.011). African descent populations were not included in the Sandholm et al. [46] study. This sign flip suggests haplotypic ancestry associated genetic architecture for this variant, such as distinct regulatory haplotypes or differential linkage patterns to other causal variants in each ancestry’s haplotypes.

#### *AKAP6* rs1885722 (chr14:32367316:T:G)

This variant is located in the A-kinase anchoring protein 6 (*AKAP6*) gene. *AKAP6* functions as a scaffold protein, forming multi-protein complexes containing protein kinase A (PKA), PDE4D3, Epac, Rap1, and ERK5. By localizing these complexes to the nuclear membrane and sarcoplasmic reticulum, *AKAP6* enables precise spatiotemporal control of cAMP-PKA signaling. Zhang et al. [47] highlight that aberrant cAMP-PKA signaling is involved in various tumors, with context-dependent roles: it can function as a tumor suppressor (via cell cycle control, apoptosis induction, and differentiation) or a tumor promoter (via enhanced migration, invasion, and metabolic reprogramming) [48]. Furthermore, Ray and Chatterjee [49] identified pleiotropy in this region affecting type 2 diabetes and prostate cancer, and Brandes, Linial, and Linial [50] reported an association with colorectal cancer. We observe protective effects for colorectal cancer in European-like ancestry haplotypes (**EUR**_SNP_ = −0.010, **EUR**_ANC_ = −0.011) and risk-increasing effects in African and Indigenous American-like ancestry haplotypes (**AFR**_SNP_ = 0.015, **AFR**_ANC_ = 0.009, **AMR**_SNP_ = 0.003, **AMR**_ANC_ = 0.003), suggesting rs1885722 may tag distinctly different nearby functional variants or different regulatory haplotypes exist in these different ancestry backgrounds, which standard, single effect SNP-only models cannot resolve.

### Coefficient maps illustrate shared and haplotypic ancestry-associated effects

A combined Manhattan-style view across phenotypes summarizes the overall sparsity pattern in Combine-S, where most ancestry-specific SNP and local-ancestry coefficients remain near zero, but a small subset of loci exhibit large ancestry-dependent magnitudes and occasional sign differences (Figure 3B). Coefficient maps for CRP show a mixture of shared SNP effects and ancestry-associated terms (Figure 4). Combine-S yields per-ancestry SNP effects that reveal flips in effect direction and ancestry-associated effect magnitudes at several loci, while modestly reducing aggregate predictive accuracy relative to Combine-R at current sample sizes (Table 1). The phased representation improves biological interpretability and discovery potential in exchange for a larger parameterization. This tradeoff should narrow as admixed sample sizes increase, as ancestry-specific effects can be estimated more precisely.

**Figure 3.**
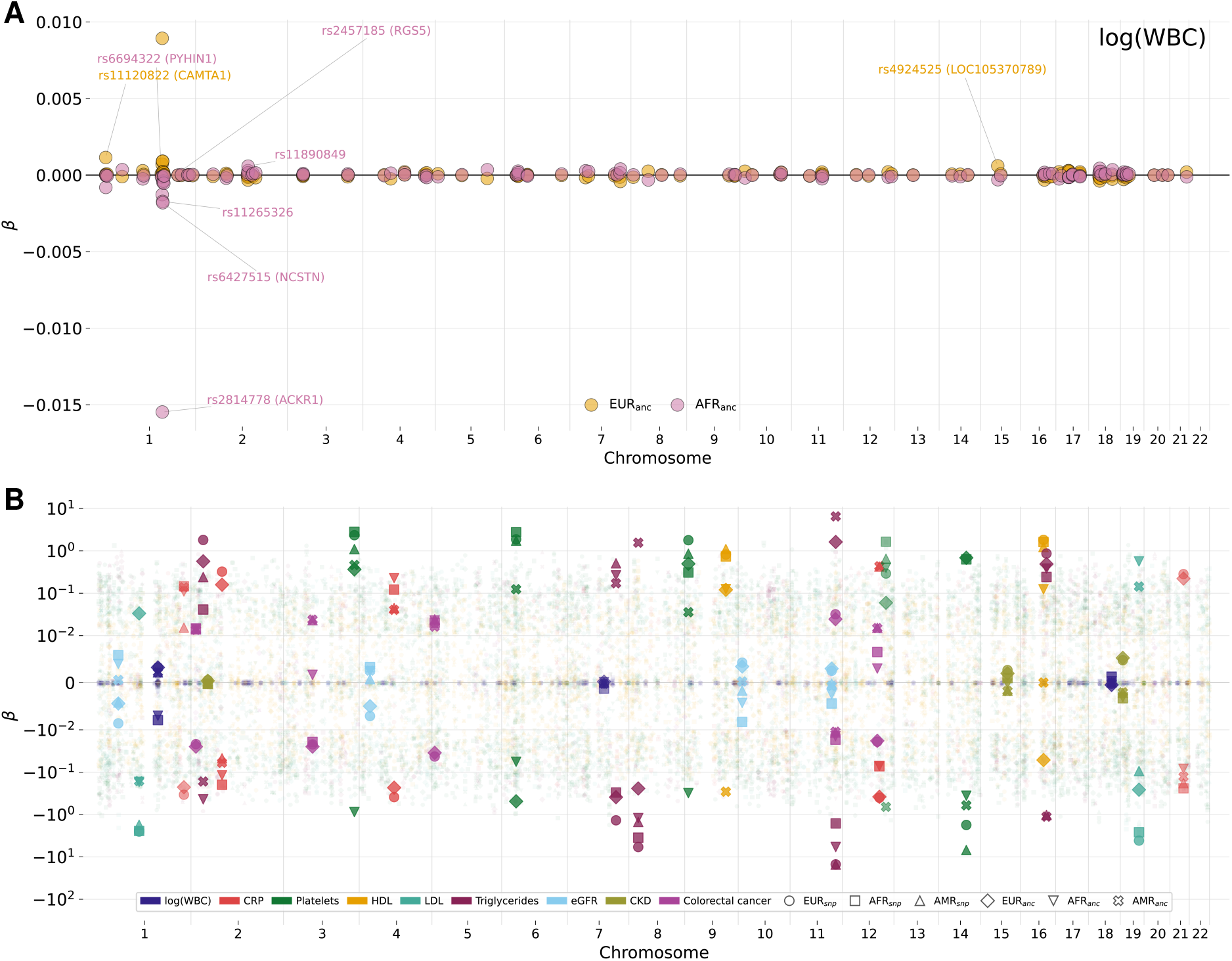
Coefficient overview for Combine-R and Combine-S. (**A**) Manhattan plot showing the EUR_ANC_ and AFR_ANC_ Combine-R coefficients for log(WBC). (**B**) Manhattan plot showing all Combine-S coefficients for all phenotypes (colors). Coefficients are plotted on a symmetric logarithmic y-axis (linear within ± 0.01). Large-magnitude coefficients are highlighted, showing strong ancestry-dependent effects (shapes).

**Figure 4.**
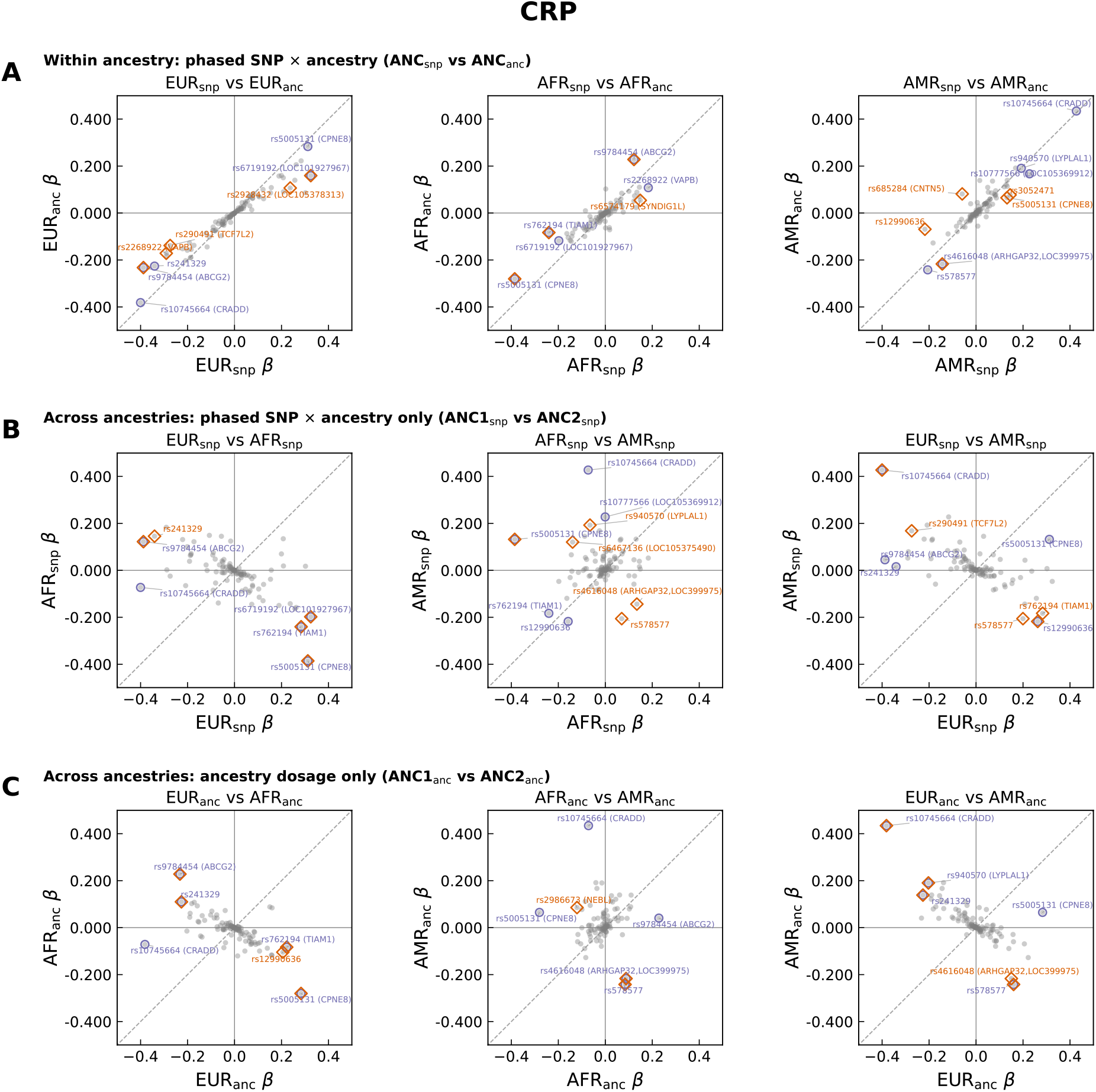
Detailed Combine-S coefficient patterns for CRP. Pairwise comparisons of estimated beta coefficients across haplotypic ancestry-associated terms are shown for variants with non-zero effects. Gray points show all variants; labels indicate variant IDs for highlighted loci. (**A**) Within-ancestry effects between phased SNP coefficients and ancestry-dosage coefficients. (**B**) SNP coefficients across ancestries. (**C**) Ancestry-dosage coefficients across ancestries. Purple highlights mark loci with the largest overall effect magnitude; orange highlights mark loci with the strongest between-axis differential effects.

### External GWAS priors improve LDL prediction in admixed cohorts

We optionally incorporate external GWAS summary statistics as univariate priors that modulate the group lasso penalties while fitting on individual-level genotypes. This keeps the design matrix and outcome unchanged and only alters the regularizer, steering sparsity toward loci supported by large external studies while keeping the final model compact. Related work has used univariate evidence to guide sparse regression via scaling and sign constraints [51]; here, GWAS evidence enters through group-specific penalty weights. The full weighting scheme and objective are described in Methods.

We demonstrate this approach for LDL cholesterol, leveraging the Global Lipids Genetics Consortium (GLGC) multi-ancestry meta-analysis [52]. We use genomic-control corrected 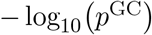 to downweight penalties at GWAS-supported loci while keeping all variants eligi-ble. LDL is also the phenotype where Combine-R is most challenged, making it a useful test case for whether external evidence can improve performance in admixed cohorts. Incorporating these priors yielded a relative increase in LDL *R*^2^ of 4.34% for Combine-S and 10.14% for Combine-R, and Combine-R with priors surpassed `snpnet` by 4.1%.

## Discussion

We have introduced a novel method, Combine, a haplotypic ancestry-aware sparse regression framework for polygenic prediction that augments SNP dosages with local ancestry and estimates joint effects using a group lasso penalty. Compared to state-of-the-art multi-ancestry summary-statistic approaches, leveraging local ancestry in 99,298 admixed All of Us participants resulted in substantial gains in prediction accuracy, with relative improvements over PRS-CSx ranging from 25.0% (CRP) to 144.2% (WBC). Furthermore, against highly optimized individual-level SNP-only models, Combine-R improved or matched predictive performance for seven of nine evaluated complex traits, while uniquely providing locus-level interpretability to disentangle shared allelic effects from ancestry-linked tagging. Finally, we demonstrated that incorporating external GWAS-derived priors further improved prediction accuracy for LDL by 10.14%. Combine-S produced per-ancestry SNP effects that increased interpretability with similar accuracy for several traits at current sample sizes. This is notable given that estimating separate effects for each ancestry necessitates having much larger number of samples, since each coefficient is fit based only on the subset of haplotypes assigned to that ancestry at that locus. Thus, with larger diverse datasets, we expect our model to increase its performance relative to simple single effect models.

Local ancestry captures variation in allele frequencies and LD structure along admixed genomes, so a single, per-variant effect is often misspecified. Combine groups each variant’s SNP and local ancestry features under a shared group penalty, letting the data decide when ancestry is informative while shrinking uninformative groups. This formulation links PRS and admixture mapping within a single estimator: Combine-R can be interpreted as learning a shared SNP effect together with ancestry-specific offsets at each locus.

Combine complements summary-statistic and association-based methods while avoiding several constraints for admixed genomes. Multi-ancestry approaches such as PRS-CSx require assigning each sample to an ancestry-matched LD panel [28], an assumption our empirical benchmarks show limits their performance in admixed cohorts compared to individual-level local-ancestry modeling. Meanwhile, local-ancestry-aware approaches like Tractor incorporate local ancestry for association testing but remain univariate for prediction [29]. Similarly, previous prediction methods designed for admixed populations utilize LAI but currently rely on preselected variants, summary statistics, or lower-dimensional ancestry models. Population-level LD panels assume a single ancestry background per reference panel, which is appropriate for single-ancestry cohorts but misaligned with the mosaic of chromosomal ancestry segments in admixed genomes; local ancestry instead lets LD structure vary along each chromosome within an individual. Combine fits directly on individual-level genotypes and local ancestry without ancestry-matched LD panels or prefiltering, accommodating arbitrary ancestry complexity. Notably, even large-scale inclusive SNP-only models have been shown to underperform on neutrophil counts due to the strong ancestry-associated effects at the *ACKR1* locus [27], a gap that our joint genotype-ancestry modeling resolves.

Combine-R, which models an unphased allele dosage plus local ancestry per variant, gave the most consistent accuracy gains, consistent with shared causal effects across ancestries for most variants and ancestry-dependent tagging through LD and MAF differences [53–55]. Combine-S estimates per-ancestry SNP effects, increasing flexibility and revealing ancestry-tagged magnitudes, but at the cost of a larger parameterization. As admixed sample sizes increase, the performance gap between Combine-S and Combine-R should narrow for traits with stronger ancestry-dependent heterogeneity. In practice, Combine-S can reveal ancestry-associated SNP differences at selected loci and cases where some ancestry-associated SNP effects are near zero but ancestry-only coefficients remain nonzero, tagging an ancestry haplotype akin to admixture mapping. These patterns are difficult to detect in SNP-only models. More generally, large SNP coefficients with near-zero ancestry-only terms suggest causal allelic drivers that do not rely on ancestry tagging, whereas the reverse pattern points to haplotypic ancestry-associated structure or linkage to one or more nearby causal, perhaps untyped variants.

We incorporate external evidence through a weighted group lasso that downweights penalties for variants supported by GWAS, while keeping all features eligible. GLGC-derived priors improved LDL prediction for Combine-R beyond both its unweighted version and the SNP-only baseline. This strategy complements meta-predictors built from univariate signals and can use multi-ancestry or ancestry-associated priors, including outputs from LAI-aware association tests such as Tractor.

The implementation scales to genome-wide fits in under 20 minutes per phenotype with millions of predictors and multiple ancestries in biobank-scale cohorts [56], as illustrated by our analysis of roughly one hundred thousand admixed participants with six ancestry categories. While Combine-R and Combine-S model significantly more features than standard SNP-only approaches, our benchmarks show that the training time does not scale linearly with the feature count; instead, efficient block-based operations allow Combine to process millions of ancestry-expanded features with per-feature costs significantly lower than the baseline. Avoiding LD matrices and summary-statistic pre-processing further simplifies the pipeline and reduces memory traffic, enabling training on the full set of typed and imputed variants rather than a preselected subset.

Within admixed individuals in the same family who share environments, but differ significantly in their mosaic of inherited haplotypes, modeling local ancestry helps separate genetic architecture from global environmental associations that might correlate with global (genome-wide averaged) genetic ancestry and which we regress out [57]. However, ancestry-dependent haplotypic coefficients can still reflect gene-by-environment or gene-by-ancestry interactions, so we recommend reporting both SNP and ancestry-only effects and auditing calibration across ancestry segments.

Higher-order PCs can capture local genomic structure and introduce collider bias rather than global population structure [58]. We therefore use global ancestry proportions as covariates, apply accuracy thresholds to LAI calls, and model local ancestry explicitly within Combine.

Several limitations remain. Performance depends on the accuracy of local ancestry inference and phasing, so errors in these could dilute ancestry-associated haplotypic effects and bias coefficients toward zero. Per-ancestry sample sizes vary along the genome, limiting power for rare segments and likely contribute to the performance gap for Combine-S. Finally, generalization to other admixture histories will require validation in additional cohorts and LAI settings.

Future work includes applying the framework to larger admixed datasets with additional ancestral representation, extending to multi-trait settings, and incorporating multi-ancestry and ancestry-associated priors in the penalty weights.

In summary, Combine integrates local ancestry into biobank-scale sparse regression for admixed genomes, producing genome-wide predictors with locus-level interpretability. Across nine phenotypes in a large admixed cohort, Combine-R achieves predictive performance comparable to a strong SNP-only lasso baseline while exposing per-variant local-ancestry coefficients that disentangle shared allelic contributions from haplotypic ancestry-linked tagging and ancestry-associated differential haplotype structure. Combine-S further enables haplotypic ancestry-associated SNP effect estimation, revealing direction and magnitude differences at selected loci. By fitting end-to-end on individual-level data and optionally incorporating external GWAS evidence through weighted group penalties, Combine provides a practical framework for ancestry-aware polygenic modeling that is both scalable and interpretable in diverse and admixed biobanks.

## Methods

### Method overview

We develop Combine, a sparse regression framework that jointly models local ancestry and SNPs using the group lasso [59]. We introduce two versions. **Combine-R** groups an unphased allele dosage with local ancestry dosages for each variant (Figure 1C). **Combine-S** uses ancestry-specific allele dosages together with local ancestry dosages, producing per-ancestry SNP effects (Figure 1D). Neither version requires LD matrices or external GWAS summary statistics. Combine returns a sparse set of genome-wide coefficients.

We fit all Combine models using `adelie` [56], which implements a path-wise block-coordinate descent solver for group lasso and related penalties in generalized linear models. At each value of the regularization parameter *λ*, the solver updates one group at a time using a Newton-type step with an adaptive one-dimensional search, so group updates converge in only a few iterations even when groups are large. The implementation combines warm starts along the *λ* path with strong-rule screening and active-set cycling, so most iterations only touch a small working set of SNP groups, which is critical for biobank-scale problems.

A key feature of `adelie` is a matrix abstraction that only requires access to inner products with columns of the design matrix. For Combine, we implement this interface using compressed representations of the SNP and local ancestry blocks, allowing matrix-vector products to operate directly on the sparse genotype and ancestry encoding without ever materializing the full *n* × *p* matrix in memory. Internally, we store SNP columns in a chunk-based sparse format that exploits the low frequency of non-reference alleles: each column is partitioned into 256-sample chunks, and only chunks containing at least one non-zero entry are stored, with 8-bit offsets indexing positions within chunks. This encoding is more compact than standard sparse formats for typical allele frequency distributions and enables cache-efficient iteration during matrix-vector products. This enables ancestry-expanded models such as Combine-R and Combine-S to be fit on hundreds of thousands of individuals and millions of variants using standard hardware, and the same infrastructure also supports SNP-only and ancestry-only PRS fits.

### Cohort characteristics and local ancestry inference

We used the All of Us v7 genotyping release and constructed a multi-ancestry reference panel by merging samples from GenomeAsia Phase 1 and 2 [60, 61], the Human Genome Diversity Project (HGDP) [62], and the 1000 Genomes Project [63]. Genetic ancestry in this reference panel was estimated with ADMIXTURE [64] in unsupervised mode with K = 6, and only individuals with at least 95% of their genome assigned to a single genetic ancestry cluster were retained as references, yielding genetic clusters interpreted as African-like (AFR, *n* = 349), Indigenous American-like (AMR, *n* = 117), East Asian-like (EAS, *n* = 340), European-like (EUR, *n* = 332), Oceanian-like (OCE, *n* = 54), and South Asian-like (SAS, *n* = 334). Reference individuals and All of Us participants were phased jointly with SHAPEIT5 (v5.1.1) using default settings [65]. We then applied Gnomix [14] to infer local ancestry along the genome, using the above reference panel and the following configuration: training proportions of 0.80 and 0.15 for the two training stages with 0.05 held out for validation, generation grid 0, 2, 4, 6, 8, 12, 16, and 24, admixture rate 1.5, window size 0.2 centimorgans, smoother span of 115 markers, context ratio 2.0, and 32 computation threads. For each genomic position, ancestry labels with posterior probability below 0.9 were treated as unassigned and set to missing. Finally, individual level global ancestry proportions were obtained by summing the local ancestry indicators for each ancestry across the genome and normalizing by the total number of loci with non missing local ancestry.

We selected the 99,298 admixed individuals from the All of Us Research program by choosing individuals with < 90% of their genome assigned to a single genetic ancestry, using a final number of 1,129,888 variants. These six global genetic ancestry proportions listed above (AFR, AMR, EAS, EUR, OCE, SAS) are later included as covariates in all prediction models, together with age and sex.

### Benchmarks

Principal components (PCs) are widely used in GWAS and individual-level data PRS models [24, 27] to adjust for population structure. However, higher-order PCs (after PC1) often capture local genomic features of interest instead of global population structure, particularly in genetically admixed populations [58]. Unless otherwise noted, all iPGS and Combine models adjust for age, sex, and the six global ancestry proportions as covariates.

To benchmark against a leading multi-ancestry summary-statistic approach, we evaluated PRS-CSx using the identical cross-validation design as the individual-level methods. Within each fold, we performed ancestry-stratified discovery GWAS in the training subset using PLINK2 [66], adjusting for age, sex, and global ancestry proportions. To ensure the distinct linkage disequilibrium patterns and effect size estimates required by PRS-CSx, individuals were assigned to a discovery GWAS stratum only if their largest global ancestry proportion exceeded 50%. Participants without a single majority ancestry were assigned to an admixed “Others” category and excluded from the ancestry-stratified training GWAS, though they were retained for validation and testing. This produced ancestry-specific summary statistics for each discovery group with sufficient sample size. PRS-CSx was then trained per fold using these ancestry-stratified summary statistics to estimate ancestry-specific posterior SNP effect sizes. We computed ancestry-specific PRS in the validation and test subsets via PLINK2 scoring. Next, we learned the optimal weights across discovery ancestries in the fold-specific validation subset. Finally, we applied these optimal weights to form a single combined PRS in the held-out test set, and evaluated its performance.

### Genotype and ancestry data formats

#### Notation

Let *n* be the number of samples, *s* the number of variants, and *A* the number of ancestries. Index individuals by *i* ∈ {1, …, *n*}, variants by *j* ∈ {1, …, *s*}, ancestries by *a* ∈ {1, …, *A*}, and haplotypes by *k* ∈ {0, 1}. Let *e*_*a*_ ∈ ℝ^*A*^ be the *a*th standard basis vector and let 1_*A*_ ∈ ℝ^*A*^ be the all-ones vector. For any block-structured matrix we write horizontal concatenation as [· | ·].

#### SNP unphased

Define the unphased genotype matrix

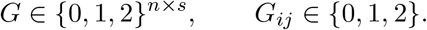

Each entry *G*_*ij*_ gives the allele dosage (number of alternate alleles) for individual *i* at variant *j*, as used in existing methods like `snpnet`.

#### SNP unphased ancestry (Combine-R) regular

For each variant *j*, define the ancestry dosage block *Y* ^(*j*)^ ∈ {0, 1, 2}^*n*×*A*^ with entries

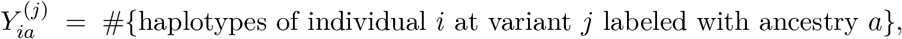

so that 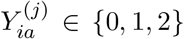 and 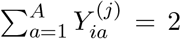 for all *i*. Let *g*^(*j*)^ := *G*_:*j*_ ∈ {0, 1, 2}^*n*^ denote the unphased genotype column. The unphased ancestry block is

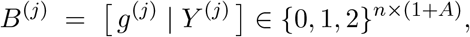

and the full matrix is

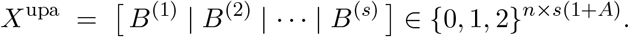

For group lasso, the 1 + *A* columns of *B*^(*j*)^ form one group.

#### SNP phased ancestry

Fix a variant *j*. For each haplotype *k* ∈ {0, 1}, define a haplotype-specific ancestry contribution

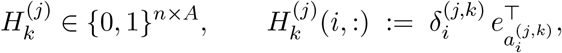

Where 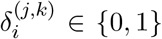 indicates whether haplotype *k* of individual *i* carries the alternate allele at variant *j*, and 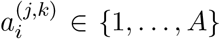 is its local-ancestry label. The phased-ancestry block at variant *j* is the sum over haplotypes,

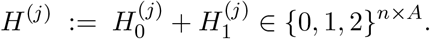

##### Example (one variant, A = 3)

Suppose genetic ancestries are labeled {1, 2, 3}. Fix a variant *j* and an individual *i*. Assume haplotype *k* = 0 carries the alternate allele and is labeled ancestry 2, while haplotype *k* = 1 does not carry the alternate allele and is labeled ancestry 3:

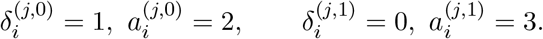

Then the per-haplotype rows are one-hot (or all-zero) vectors:

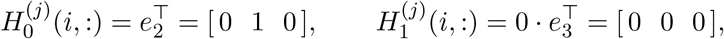

and their sum is

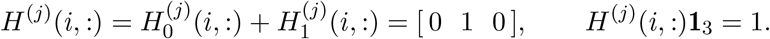

If instead both haplotypes carry the alternate allele and are labeled ancestries 2 and 3, then

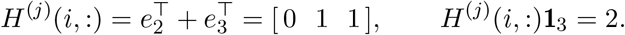

##### Interpretation

For individual *i*, the row *H*^(*j*)^(*i*, :) counts the number of alternate-allele haplotypes assigned to each ancestry. In particular,

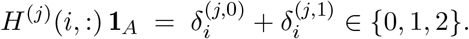

Stacking blocks across variants yields

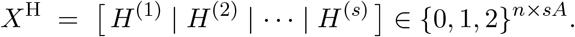

For group lasso, the *A* columns of *H*^(*j*)^ form one group.

#### SNP joint mutation and ancestry dosage (Combine-S) (specific)

For each variant *j*, combine the phased ancestry block *H*^(*j*)^ ∈ {0, 1, 2}^*n*×*A*^ and the unphased ancestry dosage block *Y* ^(*j*)^ ∈ {0, 1, 2}^*n*×*A*^ into

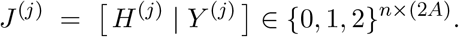

Stacking across variants yields

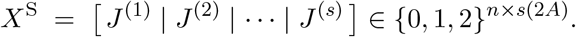

Per individual *i*, when local ancestry is assigned on both haplotypes at variant *j*,

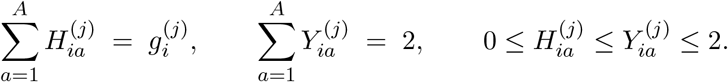

Equality in the second expression holds when both haplotypes have assigned local ancestry at variant *j. Interpretation per variant j*. Columns 1, …, *A* of *J* ^(*j*)^ give the number of alternate-allele haplotypes labeled with each ancestry, as in the phased ancestry format. Columns *A* + 1, …, 2*A* give the local ancestry dosages per ancestry, as in the unphased ancestry format. For group lasso, the 2*A* columns of *J* ^(*j*)^ form a group.

When using Combine, one can restrict to a subset *S* ⊆ {1, …, *A*} of the available ancestries, which replaces *A* by |*S*| in the ancestry-expanded formats. In our experiments, we use all 6 available ancestries for Combine-R, and subset Combine-S to EUR, AFR, and AMR due to memory limitations of the All of Us compute platform.

### External GWAS priors and weighted group lasso

We incorporate external GWAS summary statistics by introducing group-specific penalty weights in the group lasso objective (see below), without modifying the design matrix.

Let (*X, y*) denote the individual-level data with *X* ∈ ℝ^*n*×*p*^ and *y* ∈ ℝ^*n*^. Let G = {*G*_1_, …, *G*_*m*_} be a partition of the coefficient indices into variant-level groups, and let *g* ∈ {1, …, *m*} index these groups. For each variant, an external GWAS provides a *p*-value or − log_10_ (*p*). For each group *g*, we obtain a genomic-control corrected 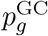 from an external GWAS across multiple cohorts. We define the GWAS score

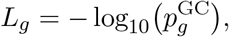

and fix a genome-wide significance level *α* = 5 × 10^−8^ with threshold *τ* = − log_10_(*α*). We construct group weights *w* = (*w*_1_, …, *w*_*m*_) ∈ ℝ^*m*^ with values in [*w*_min_, 1], where 0 < *w*_min_ < 1. Loci missing from the GWAS or not genome-wide significant (i.e., *L*_*g*_ < *τ*) receive the default weight *w*_*g*_ = 1. Let

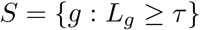

denote the set of genome-wide significant groups. Among the significant loci with *L*_*g*_ ≥ *τ*, let

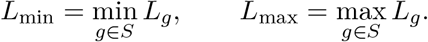

For each such locus we map the univariate signal to a smaller penalty using a linear min-max transform, For *g* ∈ *S* define

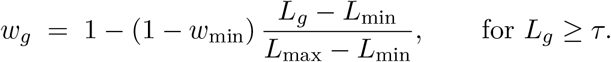

This keeps the penalty at its default for unremarkable or unobserved groups, and gradually reduces the penalty toward *w*_min_ as the external evidence strengthens.

We then fit a weighted group lasso without subsetting variants or modifying *X*. With an intercept *β*_0_ ∈ ℝ the objective is

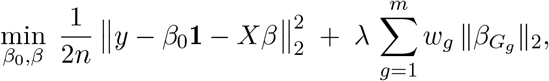

where *β*_*Gg*_ collects the coefficients in group *G*_*g*_. When no external priors are used, *w*_*g*_ = 1 for all *g* and the estimator reduces to the standard unweighted version. The hyperparameter *λ* is chosen with a held out validation set exactly as in the unweighted case.

## Acknowledgements

We gratefully acknowledge All of Us participants for their contributions, without whom this research would not have been possible. We also thank the National Institutes of Health’s All of Us Research Program for making available the participant data examined in this study.

## Data and Code Availability

Analyses were conducted in the All of Us Researcher Workbench, and access to the underlying individual level data is available to qualified researchers who enroll in the program and complete the required training and data use agreements. The group lasso matrix formats used in Combine are implemented in https://github.com/AI-sandbox/Combine.

## Supplementary Information Phenotype details

We analyzed nine phenotypes from the All of Us Research Program: log-transformed white blood cell count (log(WBC)), C-reactive protein (CRP, mg/L), platelet count, estimated glomerular filtration rate (eGFR) using the Modification of Diet in Renal Disease (MDRD) equation, triglycerides, HDL cholesterol, LDL cholesterol, colorectal cancer, and chronic kidney disease (Table S1). For all analyses we used a race neutral creatinine based eGFR, obtained by removing the Black ethnicity coefficient from the MDRD equation, motivated by evidence that the standard race adjustment systematically overestimates measured GFR [67, 68].

**Table S1.** Sample sizes for selected phenotypes across train, validation, and test splits. For quantitative traits, total sample counts are shown. For binary traits, cases/controls are reported.

| Phenotype | Type | Train | Validation | Test |
| --- | --- | --- | --- | --- |
| $\log(\text{WBC})$ | Quantitative | 20,545 | 4,453 | 4,473 |
| CRP (mg/L) | Quantitative | 9,202 | 2,029 | 2,009 |
| Platelets | Quantitative | 34,589 | 7,389 | 7,355 |
| eGFR MDRD (no race corr.) | Quantitative | 9,940 | 2,159 | 2,100 |
| Triglycerides | Quantitative | 23,009 | 5,033 | 4,940 |
| HDL | Quantitative | 22,864 | 5,008 | 4,900 |
| LDL | Quantitative | 17,468 | 3,797 | 3,735 |
| Colorectal Cancer | Binary | 399 / 68,973 | 96 / 14,919 | 77 / 14,834 |
| Chronic Kidney Disease | Binary | 4,663 / 64,709 | 1,047 / 13,968 | 994 / 13,917 |

### Benchmark

#### Extended results

Extended cross-validation results for the nine phenotypes are shown in Figure S1, and timing comparisons in Figure S2.

#### Feasibility of comparisons with existing local-ancestry methods

We did not benchmark against GAUDI [30], HAUDI [31], or SDPR_admix [32] due to structural incompatibilities with biobank-scale multi-way admixture. First, these methods are currently implemented assuming exactly two (or occasionally three) ancestry tracks. As described in Methods, we use six genetic ancestry clusters determined as the best fit via unsupervised genetic clustering to represent the All of Us cohort; reducing this complexity to fit the constraints of these more limited tools would require generating local ancestry calls for nearly 100,000 individuals using a misspecified reference panel of only two or three ancestries, yielding erroneous ancestry results. Second, GAUDI and SDPR_admix have memory constraints on genome-wide design matrices that far exceed available resources at this sample size (*N* ≈ 100*k*). While HAUDI introduces efficiency improvements over GAUDI, it remains constrained by the admixture dimensionality.

**Figure S1.**
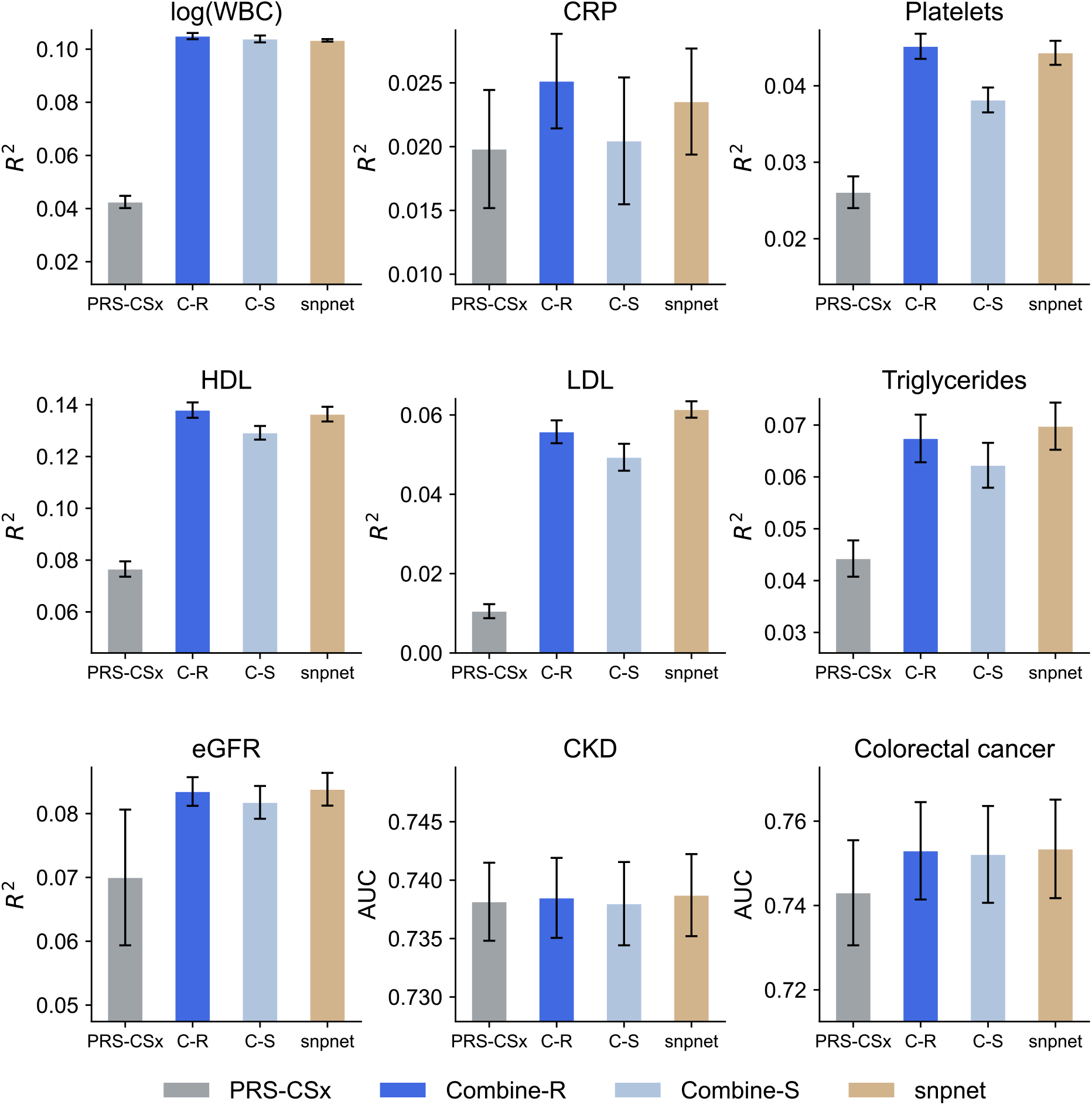
Method comparison across nine phenotypes. Bars show the mean predictive performance across 5-fold cross-validation; error bars indicate the standard error across folds.

### GLGC GWAS Summary Statistics

To inform the priors for LDL, we utilized GWAS summary statistics from the GLGC multi-ancestry meta-analysis [52]. This dataset aggregates results from 201 primary studies comprising 1,654,960 individuals across five population cohorts, which it has named: European (*N* = 1, 320, 016), East Asian (*N* = 146, 492), Admixed African or African (*N* = 99, 432), His-panic (*N* = 48, 057), and South Asian (*N* = 40, 963). The analysis evaluated associations for low-density lipoprotein cholesterol (LDL-C) covering approximately 52 million variants, which provided the genomic-control corrected 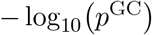 used to compute the penalty weights.

**Figure S2.**
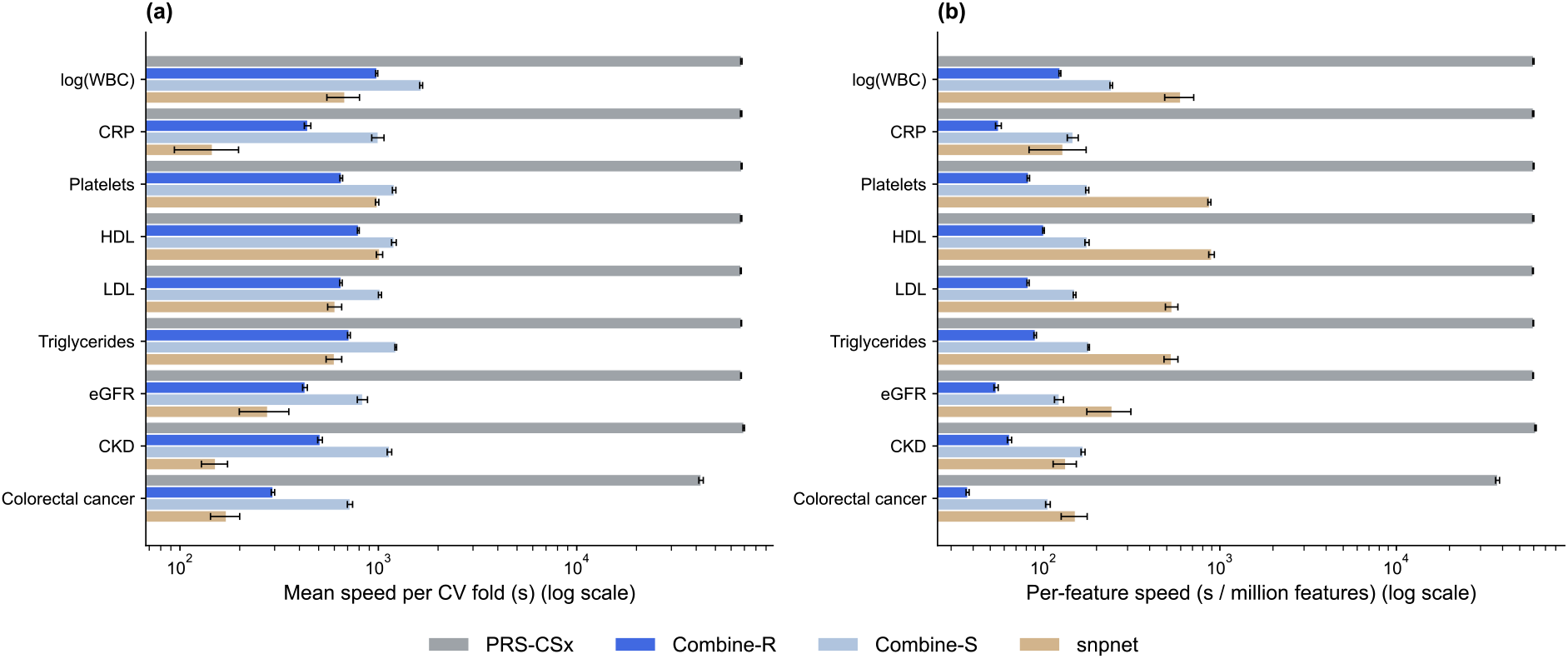
Computational efficiency. (a) Mean wall-clock time per cross-validation fold for Combine-R, Combine-S, and `snpnet` (iPGS). Despite modeling several times more features per variant, Combine’s total training time is comparable to the SNP-only baseline. (b) Normalized training time per million features. Combine-R and Combine-S are consistently more efficient per feature, leveraging compressed matrix operations to handle the expanded feature space without a proportional increase in runtime. Error bars represent the standard error across folds.

## References

1. Wray, N. R., Goddard, M. E., and Visscher, P. M. (2007). Prediction of individual genetic risk to disease from genome-wide association studies. Genome Research 17, 1520–1528.

2. Abdellaoui, A., Yengo, L., Verweij, K. J., and Visscher, P. M. (2023). 15 years of GWAS discovery: realizing the promise. The American Journal of Human Genetics 110, 179–194.

3. Bustamante, C. D., De La Vega, F. M., and Burchard, E. G. (2011). Genomics for the world. Nature 475, 163–165.

4. Dalvie, S., Koen, N., Duncan, L., Abbo, C., Akena, D., Atwoli, L., Chiliza, B., Donald, K. A., Kinyanda, E., Lochner, C., et al. (2015). Large scale genetic research on neuropsychiatric disorders in African populations is needed. eBioMedicine 2, 1259–1261.

5. Martin, A. R., Gignoux, C. R., Walters, R. K., Wojcik, G. L., Neale, B. M., Gravel, S., Daly, M. J., Bustamante, C. D., and Kenny, E. E. (2017). Human demographic history impacts genetic risk prediction across diverse populations. The American Journal of Human Genetics 100, 635–649.

6. Duncan, L., Shen, H., Gelaye, B., Meijsen, J., Ressler, K., Feldman, M., Peterson, R., and Domingue, B. (2019). Analysis of polygenic risk score usage and performance in diverse human populations. Nature Communications 10, 3328.

7. Wang, Y., Guo, J., Ni, G., Yang, J., Visscher, P. M., and Yengo, L. (2020). Theoretical and empirical quantification of the accuracy of polygenic scores in ancestry divergent populations. Nature Communications 11, 3865.

8. Ding, Y., Hou, K., Xu, Z., Pimplaskar, A., Petter, E., Boulier, K., Privé, F., Vilhjálmsson, B. J., Olde Loohuis, L. M., and Pasaniuc, B. (2023). Polygenic scoring accuracy varies across the genetic ancestry continuum. Nature 618, 774–781.

9. Moreno-Grau, S., Vernekar, M., Lopez-Pineda, A., Mas-Montserrat, D., Barrabés, M., Quinto-Cortés, C. D., Moatamed, B., Lee, M. T. M., Yu, Z., Numakura, K., et al. (2024). Polygenic risk score portability for common diseases across genetically diverse populations. Human Genomics 18, 93.

10. Kachuri, L., Chatterjee, N., Hirbo, J., Schaid, D. J., Martin, I., Kullo, I. J., Kenny, E. E., Pasaniuc, B., Polygenic Risk Methods in Diverse Populations (PRIMED) Consortium Methods Working Group, Witte, J. S., et al. (2024). Principles and methods for transferring polygenic risk scores across global populations. Nature Reviews Genetics 25, 8–25.

11. Petrovski, S. and Goldstein, D. B. (2016). Unequal representation of genetic variation across ancestry groups creates healthcare inequality in the application of precision medicine. Genome Biology 17, 157.

12. Martin, A. R., Kanai, M., Kamatani, Y., Okada, Y., Neale, B. M., and Daly, M. J. (2019). Clinical use of current polygenic risk scores may exacerbate health disparities. Nature Genetics 51, 584–591.

13. Maples, B. K., Gravel, S., Kenny, E. E., and Bustamante, C. D. (2013). RFMix: a discriminative modeling approach for rapid and robust local-ancestry inference. The American Journal of Human Genetics 93, 278–288.

14. Hilmarsson, H., Kumar, A. S., Rastogi, R., Bustamante, C. D., Montserrat, D. M., and Ioannidis, A. G. (2021). High resolution ancestry deconvolution for next generation genomic data. bioRxiv, 2021–09.

15. Skotte, L., Jørsboe, E., Korneliussen, T. S., Moltke, I., and Albrechtsen, A. (2019). Ancestry-specific association mapping in admixed populations. Genetic Epidemiology 43, 506–521.

16. Marnetto, D., Pärna, K., Läll, K., Molinaro, L., Montinaro, F., Haller, T., Metspalu, M., Mägi, R., Fischer, K., and Pagani, L. (2020). Ancestry deconvolution and partial polygenic score can improve susceptibility predictions in recently admixed individuals. Nature Communications 11, 1628.

17. Mani, A. (2017). Local Ancestry Association, Admixture Mapping, and Ongoing Challenges. Circulation: Cardiovascular Genetics 10 (2).

18. Carrot-Zhang, J., Soca-Chafre, G., Patterson, N., Thorner, A. R., Nag, A., Watson, J., Genovese, G., Rodriguez, J., Gelbard, M. K., Corrales-Rodriguez, L., et al. (2021). Genetic ancestry contributes to somatic mutations in lung cancers from admixed Latin American populations. Cancer Discovery 11, 591–598.

19. Rhead, B., Hein, D. M., Pouliot, Y., Guinney, J., De La Vega, F. M., and Sanford, N. N. (2024). Association of genetic ancestry with molecular tumor profiles in colorectal cancer. Genome Medicine 16, 99.

20. Arora, K., Tran, T. N., Kemel, Y., Mehine, M., Liu, Y. L., Nandakumar, S., Smith, S. A., Brannon, A. R., Ostrovnaya, I., Stopsack, K. H., et al. (2022). Genetic ancestry correlates with somatic differences in a real-world clinical cancer sequencing cohort. Cancer Discovery 12, 2552–2565.

21. Amuzu, S., Xie, A. X., Bai, X., Pekala, K. R., Pickersgill, N. A., Ma, D., Perea-Chamblee, T., Arora, K., Chatila, W. K., Derkach, A., et al. (2025). Meta-analysis reveals differences in somatic alterations by genetic ancestry across common cancers. Nature Genetics, 1–6.

22. Hall, M., Tach, L., and Lee, B. A. (2016). Trajectories of ethnoracial diversity in American communities, 1980–2010. Population and Development Review 42, 271.

23. Tibshirani, R. (1996). Regression shrinkage and selection via the lasso. Journal of the Royal Statistical Society Series B: Statistical Methodology 58, 267–288.

24. Qian, J., Tanigawa, Y., Du, W., Aguirre, M., Chang, C., Tibshirani, R., Rivas, M. A., and Hastie, T. (2020). A fast and scalable framework for large-scale and ultrahigh-dimensional sparse regression with application to the UK Biobank. PLOS Genetics 16, e1009141.

25. Sinnott-Armstrong, N., Tanigawa, Y., Amar, D., Mars, N., Benner, C., Aguirre, M., Venkataraman, G. R., Wainberg, M., Ollila, H. M., Kiiskinen, T., et al. (2021). Genetics of 35 blood and urine biomarkers in the UK Biobank. Nature Genetics 53, 185–194.

26. Tanigawa, Y., Qian, J., Venkataraman, G., Justesen, J. M., Li, R., Tibshirani, R., Hastie, T., and Rivas, M. A. (2022). Significant sparse polygenic risk scores across 813 traits in UK Biobank. PLOS Genetics 18, e1010105.

27. Tanigawa, Y. and Kellis, M. (2023). Power of inclusion: Enhancing polygenic prediction with admixed individuals. The American Journal of Human Genetics 110, 1888–1902.

28. Ruan, Y., Lin, Y.-F., Feng, Y.-C. A., Chen, C.-Y., Lam, M., Guo, Z., He, L., Sawa, A., Martin, A. R., et al. (2022). Improving polygenic prediction in ancestrally diverse populations. Nature Genetics 54, 573–580.

29. Atkinson, E. G., Maihofer, A. X., Kanai, M., Martin, A. R., Karczewski, K. J., Santoro, M. L., Ulirsch, J. C., Kamatani, Y., Okada, Y., Finucane, H. K., et al. (2021). Tractor uses local ancestry to enable the inclusion of admixed individuals in GWAS and to boost power. Nature Genetics 53, 195–204.

30. Sun, Q., Rowland, B. T., Chen, J., Mikhaylova, A. V., Avery, C., Peters, U., Lundin, J., Matise, T., Buyske, S., Tao, R., et al. (2024). Improving polygenic risk prediction in admixed populations by explicitly modeling ancestral-differential effects via GAUDI. Nature Communications 15, 1016.

31. Ockerman, F., Chen, B., Sun, Q., Kharitonova, E. V., Chen, W., Zhou, L. Y., Loos, R. J., Kooperberg, C., Peters, U., Haessler, J., et al. (2025). An Efficient Lasso Framework for Admixture-Aware Polygenic Scores. bioRxiv, 2025–08.

32. Zhou, G., Yolou, I., Xie, Y., and Zhao, H. (2025). Leveraging local ancestry and cross-ancestry genetic architecture to improve genetic prediction of complex traits in admixed populations. The American Journal of Human Genetics 112, 1923–1935.

33. Tsuo, K., Shi, Z., Ge, T., Mandla, R., Hou, K., Ding, Y., Pasaniuc, B., Wang, Y., and Martin, A. R. (2025). All of Us diversity and scale improve polygenic prediction contextually with greatest improvements for under-represented populations. bioRxiv, 2024–08.

34. Reich, D., Nalls, M. A., Kao, W. L., Akylbekova, E. L., Tandon, A., Patterson, N., Mullikin, J., Hsueh, W.-C., Cheng, C.-Y., Coresh, J., et al. (2009). Reduced neutrophil count in people of African descent is due to a regulatory variant in the Duffy antigen receptor for chemokines gene. PLOS Genetics 5, e1000360.

35. The All of Us Research Program Genomics Investigators (2024). Genomic data in the All of Us Research Program. Nature 627, 340.

36. Van Driest, S. L., Abul-Husn, N. S., Glessner, J. T., Bastarache, L., Nirenberg, S., Schildcrout, J. S., Eswarappa, M. S., Belbin, G. M., Shaffer, C. M., Mentch, F., et al. (2021). Association between a common, benign genotype and unnecessary bone marrow biopsies among African American patients. JAMA Internal Medicine 181, 1100–1105.

37. Chung, C. P., Karakoc, G., Liu, G., Gamboa, J. L., Mosley, J. D., Cox, N. J., Stein, C. M., and Kawai, V. (2022). Ancestry, ACKR1 and leucopenia in patients with systemic lupus erythematosus. Lupus Science & Medicine 9, e000790.

38. Pirim, D., Wang, X., Niemsiri, V., Radwan, Z. H., Bunker, C. H., Hokanson, J. E., Hamman, R. F., Barmada, M. M., Demirci, F. Y., and Kamboh, M. I. (2016). Resequencing of the CETP gene in American whites and African blacks: Association of rare and common variants with HDL-cholesterol levels. Metabolism 65, 36–47.

39. Chang, M.-h., Ned, R. M., Hong, Y., Yesupriya, A., Yang, Q., Liu, T., Janssens, A. C. J., and Dowling, N. F. (2011). Racial/ethnic variation in the association of lipid-related genetic variants with blood lipids in the US adult population. Circulation: Cardiovascular Genetics 4, 523–533.

40. Salari, N., Darvishi, N., Mansouri, K., Ghasemi, H., Hosseinian-Far, M., Darvishi, F., and Mohammadi, M. (2021). Association between PNPLA3 rs738409 polymorphism and nonalcoholic fatty liver disease: a systematic review and meta-analysis. BMC Endocrine Disorders 21, 125.

41. Cox, A. J., Wing, M. R., Carr, J. J., Hightower, R. C., Smith, S. C., Xu, J., Wagenknecht, L. E., Bowden, D. W., and Freedman, B. I. (2011). Association of PNPLA3 SNP rs738409 with liver density in African Americans with type 2 diabetes mellitus. Diabetes & Metabolism 37, 452–455.

42. Liska, D., Dufour, S., Zern, T. L., Taksali, S., Calí, A. M., Dziura, J., Shulman, G. I., Pierpont, B. M., and Caprio, S. (2007). Interethnic differences in muscle, liver and abdominal fat partitioning in obese adolescents. PLOS One 2, e569.

43. Guerrero, R., Vega, G. L., Grundy, S. M., and Browning, J. D. (2009). Ethnic differences in hepatic steatosis: an insulin resistance paradox? Hepatology 49, 791–801.

44. Yu, S. S., Castillo, D. C., Courville, A. B., and Sumner, A. E. (2012). The Triglyceride Paradox in People of African Descent. Metabolic Syndrome and Related Disorders 10, 77–82.

45. Uhlén, M., Fagerberg, L., Hallström, B. M., Lindskog, C., Oksvold, P., Mardinoglu, A., Sivertsson, Å., Kampf, C., Sjöstedt, E., Asplund, A., et al. (2015). Tissue-based map of the human proteome. Science 347, 1260419.

46. Sandholm, N., Cole, J. B., Nair, V., Sheng, X., Liu, H., Ahlqvist, E., Van Zuydam, N., Dahlström, E. H., Fermin, D., Smyth, L. J., et al. (2022). Genome-wide meta-analysis and omics integration identifies novel genes associated with diabetic kidney disease. Diabetologia 65, 1495–1509.

47. Zhang, H., Kong, Q., Wang, J., Jiang, Y., and Hua, H. (2020). Complex roles of cAMP– PKA–CREB signaling in cancer. Experimental Hematology & Oncology 9, 32.

48. Rosenthal, K. J., Gordan, J. D., and Scott, J. D. (2024). Protein kinase A and local signaling in cancer. Biochemical Journal 481, 1659–1677.

49. Ray, D. and Chatterjee, N. (2020). A powerful method for pleiotropic analysis under composite null hypothesis identifies novel shared loci between Type 2 Diabetes and Prostate Cancer. PLOS Genetics 16, e1009218.

50. Brandes, N., Linial, N., and Linial, M. (2021). Genetic association studies of alterations in protein function expose recessive effects on cancer predisposition. Scientific Reports 11, 14901.

51. Chatterjee, S., Hastie, T., and Tibshirani, R. (2025). Univariate-guided sparse regression. arXiv preprint arXiv:2501.18360.

52. Graham, S. E., Clarke, S. L., Wu, K.-H. H., Kanoni, S., Zajac, G. J., Ramdas, S., Surakka, I., Ntalla, I., Vedantam, S., Winkler, T. W., et al. (2021). The power of genetic diversity in genome-wide association studies of lipids. Nature 600, 675–679.

53. Hou, K., Ding, Y., Xu, Z., Wu, Y., Bhattacharya, A., Mester, R., Belbin, G. M., Buyske, S., Conti, D. V., Darst, B. F., et al. (2023). Causal effects on complex traits are similar for common variants across segments of different continental ancestries within admixed individuals. Nature Genetics 55, 549–558.

54. Carlson, C. S., Matise, T. C., North, K. E., Haiman, C. A., Fesinmeyer, M. D., Buyske, S., Schumacher, F. R., Peters, U., Franceschini, N., Ritchie, M. D., et al. (2013). Generalization and dilution of association results from European GWAS in populations of non-European ancestry: the PAGE study. PLOS Biology 11, e1001661.

55. Mägi, R., Horikoshi, M., Sofer, T., Mahajan, A., Kitajima, H., Franceschini, N., McCarthy, M. I., COGENT-Kidney Consortium, T.-G. C., and Morris, A. P. (2017). Trans-ethnic meta-regression of genome-wide association studies accounting for ancestry increases power for discovery and improves fine-mapping resolution. Human Molecular Genetics 26, 3639–3650.

56. Yang, J. and Hastie, T. (2024). A fast and scalable pathwise-solver for group lasso and elastic net penalized regression via block-coordinate descent. arXiv preprint arXiv:2405.08631.

57. Atkinson, E. G. (2023). Estimation of cross-ancestry genetic correlations within ancestry tracts of admixed samples. Nature Genetics 55, 527–529.

58. Grinde, K. E., Browning, B. L., Reiner, A. P., Thornton, T. A., and Browning, S. R. (2024). Adjusting for principal components can induce collider bias in genome-wide association studies. PLOS Genetics 20, e1011242.

59. Yuan, M. and Lin, Y. (2006). Model selection and estimation in regression with grouped variables. Journal of the Royal Statistical Society Series B: Statistical Methodology 68, 49–67.

60. The GenomeAsia 100K Project enables genetic discoveries across Asia (2019). Nature 576, 106–111.

61. Wall, J. D., Sathirapongsasuti, J. F., Gupta, R., Rasheed, A., Venkatesan, R., Belsare, S., Menon, R., Phalke, S., Mittal, A., Fang, J., et al. (2023). South Asian medical cohorts reveal strong founder effects and high rates of homozygosity. Nature Communications 14, 3377.

62. Bergström, A., McCarthy, S. A., Hui, R., Almarri, M. A., Ayub, Q., Danecek, P., Chen, Y., Felkel, S., Hallast, P., Kamm, J., et al. (2020). Insights into human genetic variation and population history from 929 diverse genomes. Science 367, eaay5012.

63. The 1000 Genomes Project Consortium (2015). A global reference for human genetic variation. Nature 526, 68–74.

64. Alexander, D. H., Novembre, J., and Lange, K. (2009). Fast model-based estimation of ancestry in unrelated individuals. Genome Research 19, 1655–1664.

65. Hofmeister, R. J., Ribeiro, D. M., Rubinacci, S., and Delaneau, O. (2023). Accurate rare variant phasing of whole-genome and whole-exome sequencing data in the UK Biobank. Nature Genetics 55, 1243–1249.

66. Chang, C. C., Chow, C. C., Tellier, L. C., Vattikuti, S., Purcell, S. M., and Lee, J. J. (2015). Second-generation PLINK: rising to the challenge of larger and richer datasets. GigaScience 4, s13742–015.

67. Gama, R. M., Clery, A., Griffiths, K., Heraghty, N., Peters, A. M., Palmer, K., Kibble, H., Vincent, R. P., Sharpe, C. C., Cairns, H., et al. (2021). Estimated glomerular filtration rate equations in people of self-reported black ethnicity in the United Kingdom: inappropriate adjustment for ethnicity may lead to reduced access to care. PLOS ONE 16, e0255869.

68. Inker, L. A., Eneanya, N. D., Coresh, J., Tighiouart, H., Wang, D., Sang, Y., Crews, D. C., Doria, A., Estrella, M. M., Froissart, M., et al. (2021). New creatinine-and cystatin C–based equations to estimate GFR without race. New England Journal of Medicine 385, 1737–1749.

